# Enriching the Human Stool Microeukaryotes for Shotgun Sequencing

**DOI:** 10.64898/2026.06.24.734237

**Authors:** Ezgi Özkurt, David Schneider, Steve A. James, Isabelle Hautefort, Jennifer Ahn-Jarvis, Darren Heavens, Manuel Banzhaf, Antonietta Hayhoe, Falk Hildebrand

## Abstract

The human gut microbiome harbours a diverse community of microeukaryotes, predominantly fungi, which may potentially play important roles in gut ecology and homeostasis. Despite their potential, the study of gut microeukaryotes has been hampered by the limited sensitivity of standard sequencing approaches, which struggle to capture DNA from low-abundance microorganisms against the overwhelming background of bacterial biomass. To address this, we developed a method to selectively enrich for microeukaryotic cells in human faecal samples by depleting bacterial cells prior to metagenomic sequencing.

Through systematic comparison and optimisation at each processing step, we established a robust standard operating procedure (SOP) for microeukaryotic cell enrichment. By benchmarking this SOP across eight human faecal samples with three technical replicates each, we showed that it consistently increased microeukaryote representation in metagenomic libraries, greater microeukaryotic taxonomic diversity, and a reduced proportion of unclassified taxa. Together, these improvements enabled substantially deeper characterisation of the microeukaryotic fraction of the human gut microbiome.

---

The human gut microbiome harbours a diverse community of microeukaryotes including fungi, protists, and unicellular algae. Despite constituting only a small fraction of the total microbial biomass, microeukaryotes are increasingly recognised as important contributors to gut ecology and homeostasis (Li et al. 2019), (Zhang et al. 2022). Among these, fungi are the most extensively studied, accounting for approximately 0.01–2% of total gut microbiota composition (Auchtung et al. 2018), (Doron et al. 2021), (Auchtung et al. 2022). Fungi were detected in over 98% of all faecal samples analysed by the Human Microbiome Project (HMP) (Nash et al. 2017). Non-fungal microeukaryotes — such as protists including protozoan species — are also consistently detected in faecal samples and may exert significant influence on microbial community structure and host immunity (Chabé et al. 2017).

Despite their potential significance for gut health, research on gut microeukaryotes has been limited by technical constraints of conventional sequencing approaches. Current methods lack the sensitivity to detect DNA from low-abundance organisms in the presence of overwhelmingly dominant bacterial biomass. Microeukaryotes differ from bacteria in several key structural features, including larger cell size (Underhill and Iliev 2014), rigid cell walls, and distinct cell wall composition (Rintarhat et al. 2024). By leveraging these physical differences, we developed and optimised a method to selectively separate microeukaryotic cells from the dominant bacterial fraction, thereby enriching microeukaryotic DNA in human faecal samples prior to metagenomic sequencing (see **Figure 1** and **Supplementary Text** for further details).

**Figure 1:**
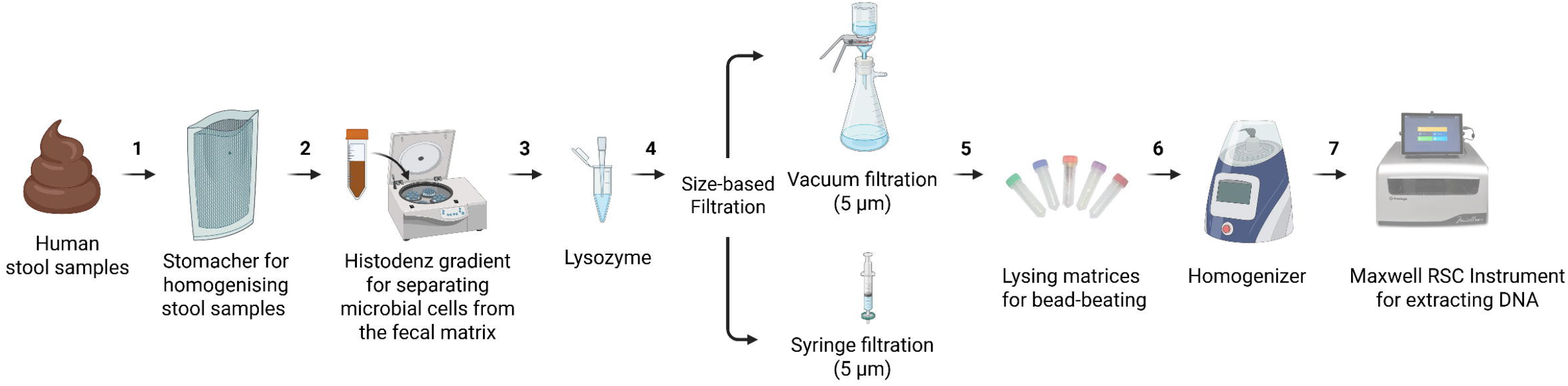
Overview of the Optimized Steps in the Protocol for Fungal Enrichment in Human Faecal Samples and Mock Microbial Communities. Faecal samples are (1) homogenized using a stomacher machine in a stomacher bag with cell strainers to remove particles larger than 5 mm. The resulting faecal suspension is applied to (2) a Histodenz density gradient to separate microbial cells from the faecal matrix. The recovered microbial suspension is treated (3) with lysozyme to lyse bacterial cells. The microbial suspensions are then (4) filtered using either a vacuum filtration unit with a cellulose nitrate membrane with 5 µm pores or a syringe filter unit with 5 µm pores. Microbial cells or membranes are (5) homogenized in a lysing matrix (6) using a FastPrep®-24 Instrument, (7) followed by DNA extraction. Özkurt, E. (2026), https://BioRender.com.

To optimise and benchmark our protocol, we first collected and processed eight faecal samples from seven volunteers. Faecal samples were collected as part of the PEARL-AGE study cohort at Quadram Institute Bioscience (ClinicalTrials.gov: NCT05346016; IRAS ID: 305505) (See **Supplementary Text** for further details). The samples were obtained from healthy donors (two female and five male; aged 10–72 years) who had not taken antibiotics within the three months preceding donation and reported no history of gastrointestinal disease at the time of donation. One of the samples was excluded from downstream analysis due to insufficient total SSU (small subunit) reads obtained after sequencing.

Six samples were used for benchmarking all protocol steps and two samples for benchmarking the cell lysis matrices. Each sample was processed in three technical replicates. Given that microeukaryotes typically occur at very low abundance in human stool, the protocol was designed to specifically retain microeukaryotic cells while removing as many bacterial cells as possible. To assess protocol performance, samples were collected after each step, from which DNA was extracted and sequenced to determine at which stage the greatest fold enrichment of microeukaryote DNA occurred.

A key challenge in developing this protocol was to efficiently separate microbial cells from the faecal matrix to enable subsequent enrichment of microeukaryotic cells relative to bacterial cells. To achieve this, we first homogenized faecal samples with a NaCl solution in a stomacher bag fitted with a 5 mm strainer, which removed larger faecal particles from the suspension. We then employed single-density gradients prepared with Histodenz, a nonionic density gradient medium, used here to retain the faecal matrix while allowing the microbial fraction to remain above the gradient.

As anticipated, the stomacher and Histodenz steps generated minimal or no enrichment (x□ = 3 ± 2.8 and x□ = 0.4 ± 0.65, respectively) **(Figure 2A)**. Enrichment became detectable after lysozyme treatment (x□ = 2.1 ± 3.3), but the major increase occurred following filtration. Because fungal cells, taken here as representative microeukaryotes, are markedly larger (∼5 μm diameter) than bacterial cells (∼1 μm) (Underhill and Iliev 2014), we used size-based filtration to separate them. Two approaches were tested: vacuum filtration using a 5 μm Cytiva AE98 cellulose nitrate membrane, and syringe filtration using a 5 μm Millex-SV filter unit. The highest mean fold enrichment was observed when syringe filtration was combined with Lysing Matrix D (x□ = 427.3 ± 802.6), followed by vacuum filtration with Lysing Matrix E (x□ = 344.3 ± 583.1) and syringe filtration with Lysing Matrix E (x□ = 294.5 ± 428.1).

**Figure 2:**
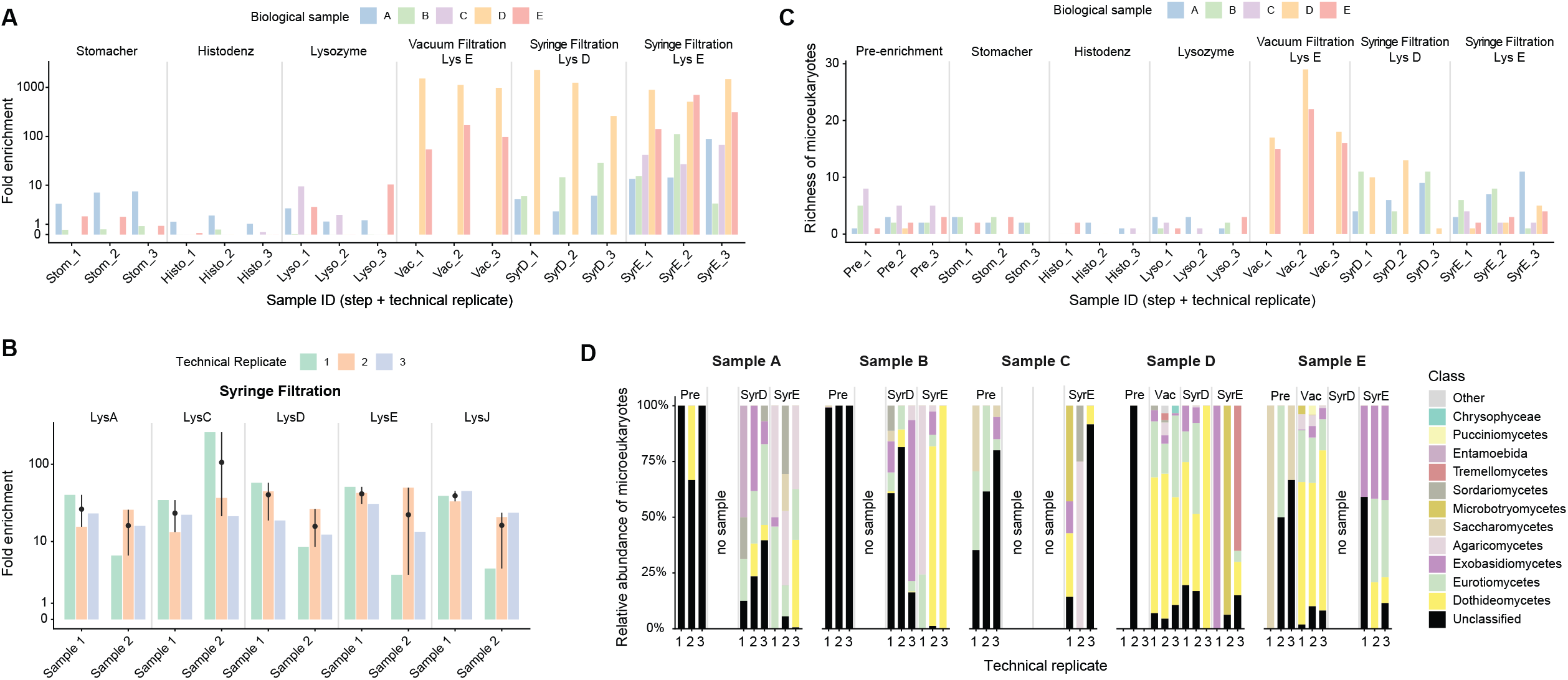
Benchmarking of each step of the protocol in human faecal samples. **A)** Fold enrichment of microeukaryotes relative to bacteria across protocol steps for each sample and technical replicate. Fold enrichment was calculated after subsampling at different protocol steps using faecal samples from six volunteers processed in three technical replicates, except for vacuum filtration and syringe filtration followed by Lysing Matrix D, for which samples from two and three volunteers were processed, respectively. **B)** Fold enrichment obtained using five different lysing matrices to process two additional human faecal samples following syringe filtration, each analysed in three technical replicates. Error bars indicate the deviation of each technical replicate from the mean of the three replicates, and the dot within each bar represents the mean fold enrichment across technical replicates. **C)** Microeukaryotic richness in samples before enrichment and after each step of the enrichment protocol. Richness (i.e. observed number) of microeukaryotes at genus level was calculated for each technical replicate of each sample after subsampling at different steps of the protocol. **D)** Relative composition of microeukaryotes in samples before enrichment and after each step of the enrichment protocol. Relative composition of microeukaryotes at class level was reported for each technical replicate of each sample after subsampling at different steps of the protocol. The top 12 classes with the highest relative abundances are shown in the figure, while all remaining classes are grouped under “Other.”

The protocol successfully depleted bacterial cells and enriched for microeukaryotes across all tested samples, with mean enrichment values for all configurations exceeding the initial target of 100-fold enrichment (x□ = 341.5 ± 471.4). However, enrichment efficiency varied substantially – ranged from 0.64 (i.e. no enrichment) to 1,741.5× – likely reflecting both biological heterogeneity in microeukaryotic loads across individuals and technical variability between replicates.

In addition, we evaluated whether lysing matrices other than D or E significantly affected fungal enrichment in two additional faecal samples **(Figure 2B)**. To this end, we compared five lysing matrices: A, C, D, E, and J (MP Biomedicals). The same enrichment protocol was applied using syringe filtration, with only the lysing matrix varied. Each matrix contained a distinct combination of bead sizes and materials, allowing us to compare conditions that maximised microeukaryotes’ cell lysis while minimising bacterial cell disruption. In faecal samples, Lysing Matrix A produced slightly lower microeukaryote-to-bacterial DNA ratios than the other matrices tested, whereas the remaining matrices showed broadly comparable ratios. Based on these results, we concluded that the use of any tested matrix other than A would be unlikely to substantially affect the microeukaryotic-to-bacterial DNA ratio.

After application of the full enrichment protocol **(Figure 1)**, microeukaryotic richness in faecal samples, measured as the observed number of genera, increased significantly **(Figure 2C)**. Before enrichment, an average of only 2.46 microeukaryotic genera were detected in faecal samples, ranging from no detected genera to eight genera **(Figure 2C)**. As expected, the stomacher, Histodenz gradient, and lysozyme steps alone did not markedly increase detected microeukaryotic richness, with mean richness values of 2.13, 0.36, and 0.87, respectively. Vacuum filtration, tested only in samples from two volunteers, significantly increased the richness of detected microeukaryotes with a mean richness of 19.3 genera and a range of 15–28 genera. When syringe filtration was used instead of vacuum filtration, microeukaryotic richness was generally higher than pre-enrichment levels but lower compared to vacuum filtration. Using syringe filtration, mean richness values increased to 7.66 with Lysing Matrix D and 4.06 with Lysing Matrix E, with ranges of 1–13 and 1–11 genera, respectively. Overall, the full enrichment protocol significantly increased the richness of detected microeukaryotes; however vacuum filtration resulted in significantly higher richness of taxa than syringe filtering.

Application of our protocol also significantly reduced the proportion of unclassified microeukaryotic taxa in faecal samples **(Figure 2D)**. In pre-enrichment samples, most taxa were unclassified at the class level, ranging from 33.3% to 99.7%. Following enrichment, a greater proportion of taxa could be assigned to known eukaryotic classes. Unclassified taxa decreased to 6.7%–7.3% after vacuum filtration with Lysing Matrix E. With syringe filtration, unclassified taxa accounted for 12.1%–52.8% after lysis with Lysing Matrix D and 0.4%–35.3% after lysis with Lysing Matrix E.

Overall, our protocol, using either vacuum or syringe filtration, successfully depleted bacterial DNA and thereby increased microeukaryotic representation in metagenomic sequencing. However, vacuum filtration yielded a higher taxonomic richness and a lower proportion of unclassified taxa than syringe filtration.

This protocol also has several limitations: biological variation in microeukaryote starting abundance, which influences enrichment efficiency; technical variability in fold-enrichment between replicates; and limited parallel throughput owing to the protocol’s length.

Samples with high bacterial loads are likely to benefit most, as bacterial depletion was highly effective without substantial loss of microeukaryotes. Samples with less complex inorganic matrices may be able to enter the protocol directly at step three. We anticipate that this protocol will also be valuable for enriching microeukaryotes from other complex microbial communities, particularly those in which microeukaryotes are more abundant and the sample matrix is less structurally complex than human stool, such as plant- or water-associated microbiomes.

## ACKNOWLEDGEMENTS

We thank the Human Studies Team at Quadram Institute Bioscience for their assistance with ethical approvals and the Clinical Research Facility with sample collection for the PEARL-AGE study, and gratefully acknowledge all participants for their contribution. We also thank E Yabuuchi, The Lister Institute, F W Hickamn-Brenner, O Lysenko, and F Griffith for depositing the bacterial strains used in this study with the NCTC and making them available to the scientific community.

## STUDY FUNDING

This work was funded by the European Research Council H2020 StG (ERC-StG-948219, EPYC) and the BBSRC Institute Strategic Programme Food Microbiome and Health (BB/X011054/1).

## CONFLICTS OF INTEREST

F.H. declares a conflict of interest as a Scientific Advisory Board member of BioMatz. All other authors declare no conflicts of interest.

## DATA AVAILABILITY

The datasets generated during and/or analysed during the current study are available in the ENA repository, PRJXXXXX.

## CODE AVAILABILITY

The data files and script necessary to reproduce the statistical analysis are provided at https://github.com/ozkurt/XXXX.

## AUTHOR CONTRIBUTIONS

**EÖ:** Conceptualization, Methodology, Validation, Formal analysis, Investigation, Resources, Writing-original draft, Writing - Review & Editing, Visualization, Supervision, Project Administration

**DS:** Methodology, Validation, Formal analysis, Investigation, Data curation, Writing – Original draft, Writing – Review & Editing, Visualization

**SAJ:** Resources, Writing - Review & Editing

**IH:** Methodology

**JAJ:** Resources

**DH:** Methodology, Resources

**MB**: Methodology, Writing - Review & Editing

**AH:** Writing - Review & Editing, Funding

**FH:** Conceptualization, Methodology, Software, Data Curation, Writing - Review & Editing, Supervision, Project administration, Funding acquisition

## Supplementary Figures

**Supplementary Figure 1.**
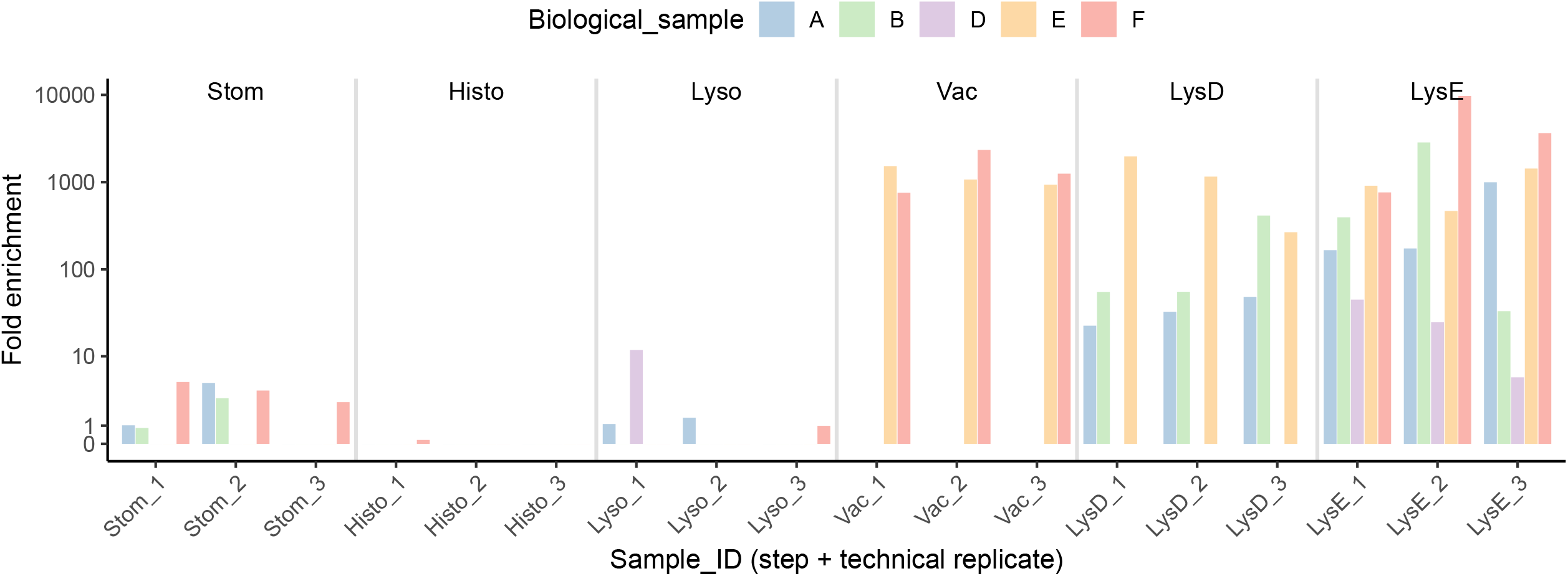
Fold enrichment of fungi relative to bacteria across protocol steps for each human faecal sample and technical replicate. Only taxa belonging to the phyla Ascomycota, Basidiomycota, and Chytridiomycota were included in the calculations.

**Suppl Figure 2:**
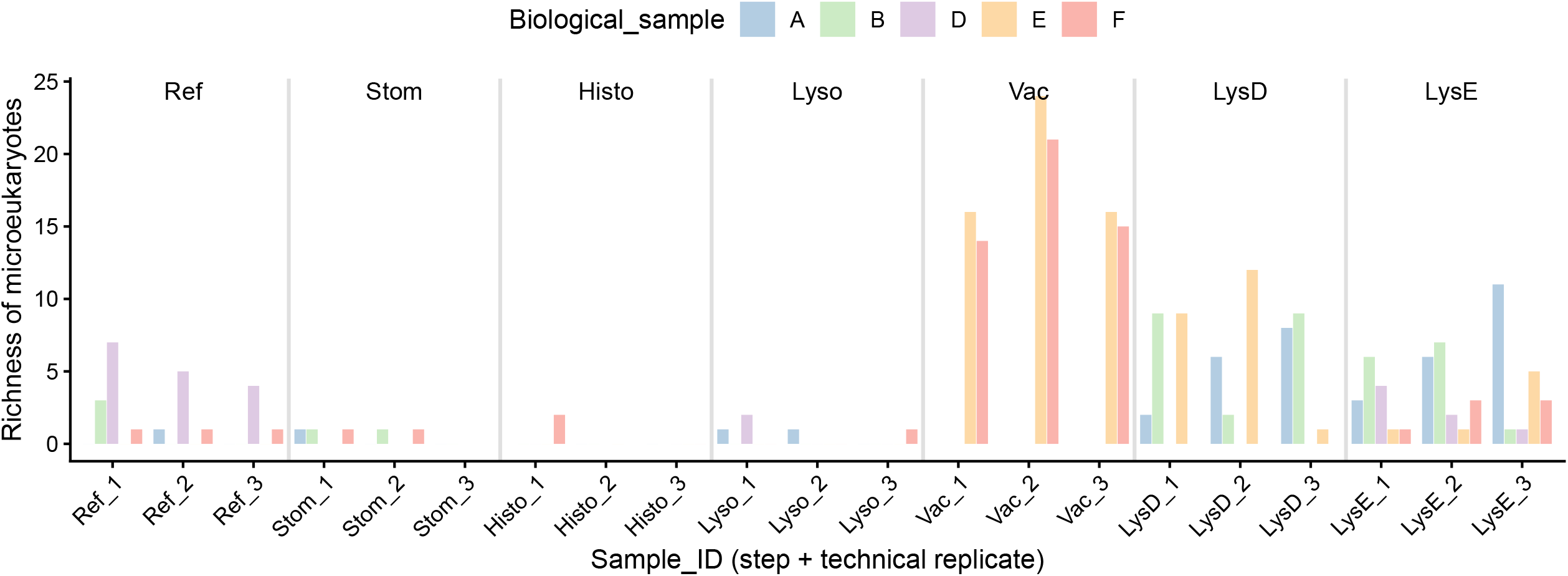
Fungal richness calculated at the genus level in human faecal samples before enrichment and after each step of the enrichment protocol. Only taxa belonging to the phyla Ascomycota, Basidiomycota, and Chytridiomycota were included in the calculations.

**Suppl Figure 3:**
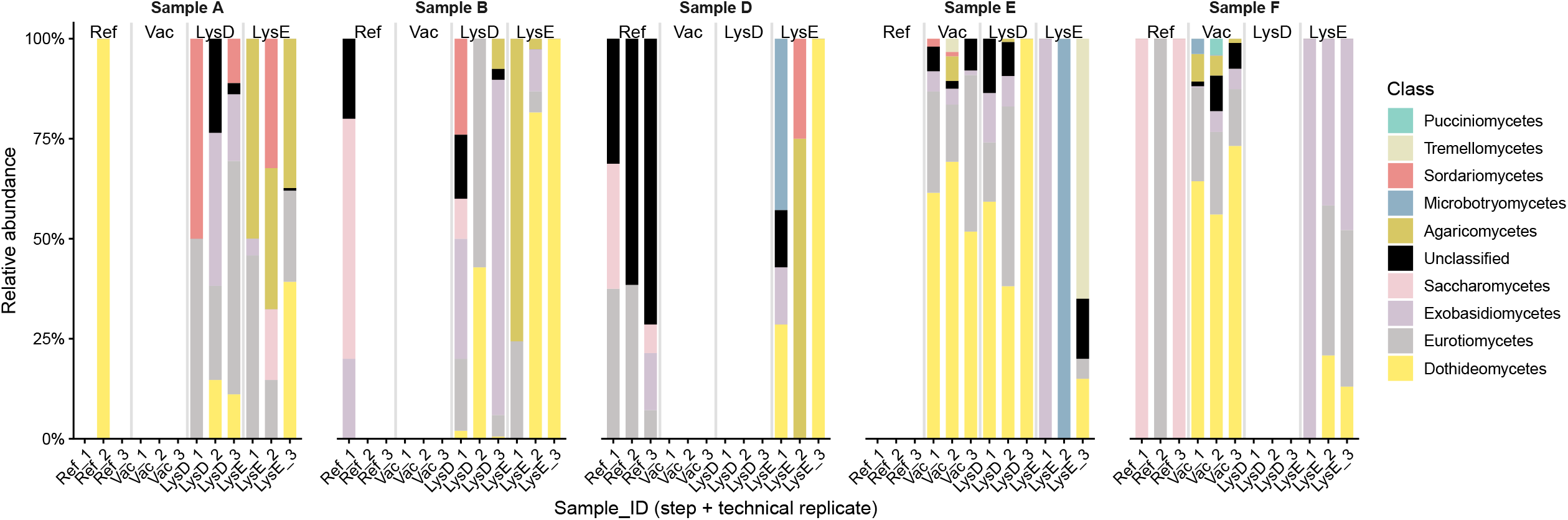
Relative composition of fungi in human faecal samples before enrichment and after each step of the enrichment protocol. Only taxa belonging to the phyla Ascomycota, Basidiomycota, and Chytridiomycota were included in the calculations.

## REFERENCES

Auchtung, Thomas A., Tatiana Y. Fofanova, Christopher J. Stewart, et al. 2018. “Investigating Colonization of the Healthy Adult Gastrointestinal Tract by Fungi.” mSphere 3 (2). 10.1128/mSphere.00092-18.

Auchtung, Thomas A., Christopher J. Stewart, Daniel P. Smith, et al. 2022. “Temporal Changes in Gastrointestinal Fungi and the Risk of Autoimmunity during Early Childhood: The TEDDY Study.” Nature Communications 13 (1): 3151.

Bahram, Mohammad, Tarquin Netherway, Clémence Frioux, et al. 2021. “Metagenomic Assessment of the Global Diversity and Distribution of Bacteria and Fungi.” Environmental Microbiology 23 (1): 316–326.

Chabé, Magali, Ana Lokmer, and Laure Ségurel. 2017. “Gut Protozoa: Friends or Foes of the Human Gut Microbiota?” Trends in Parasitology 33 (12): 925–934.

Doron, Itai, Irina Leonardi, Xin V. Li, et al. 2021. “Human Gut Mycobiota Tune Immunity via CARD9-Dependent Induction of Anti-Fungal IgG Antibodies.” Cell 184 (4): 1017–1031.e14.

Frioux, Clémence, Rebecca Ansorge, Ezgi Özkurt, et al. 2023. “Enterosignatures Define Common Bacterial Guilds in the Human Gut Microbiome.” Cell Host & Microbe 31 (7): 1111–1125.e6.

Li, Xin V., Irina Leonardi, and Iliyan D. Iliev. 2019. “Gut Mycobiota in Immunity and Inflammatory Disease.” Immunity 50 (6): 1365–1379.

Nash, Andrea K., Thomas A. Auchtung, Matthew C. Wong, et al. 2017. “The Gut Mycobiome of the Human Microbiome Project Healthy Cohort.” Microbiome 5 (1): 153.

Nnadozie, Chika F., Johnson Lin, and Roshini Govinden. 2015. “Selective Isolation of Bacteria for Metagenomic Analysis: Impact of Membrane Characteristics on Bacterial Filterability.” Biotechnology Progress 31 (4): 853–866.

Özkurt, Ezgi, Joachim Fritscher, Nicola Soranzo, et al. 2022. “LotuS2: An Ultrafast and Highly Accurate Tool for Amplicon Sequencing Analysis.” Microbiome 10 (1): 176.

Rintarhat, Piyapat, Yong-Joon Cho, Hong Koh, et al. 2024. “Assessment of DNA Extraction Methods for Human Gut Mycobiome Analysis.” Royal Society Open Science 11 (1): 231129.

Underhill, David M., and Iliyan D. Iliev. 2014. “The Mycobiota: Interactions between Commensal Fungi and the Host Immune System.” Nature Reviews. Immunology 14 (6): 405–416.

Zhang, Fen, Dominik Aschenbrenner, Ji Youn Yoo, and Tao Zuo. 2022. “The Gut Mycobiome in Health, Disease, and Clinical Applications in Association with the Gut Bacterial Microbiome Assembly.” The Lancet. Microbe 3 (12): e969–e983.

