## Supplementary Text for "Enriching the Human Stool Microeukaryotes for Shotgun Sequencing"

**Supplementary Material:**

**Sample collections:**

All participants provided written informed consent prior to sample collection, and all procedures involving human biological material were conducted in accordance with the Human Tissue Act 2004 (Human Tissue Authority). All procedures involving human biological material were conducted in accordance with relevant institutional guidelines and regulations and complied with the principles outlined in the Declaration of Helsinki. Samples were self-collected by participants, transferred to clean collection bags, and immediately placed in cool boxes with frozen cold packs. Samples were transported to the laboratory under cold chain conditions and stored at −80°C until analysis.

**Overall protocol:**

One gram of stool was mixed with 10 mL of 0.9% NaCl in a stomacher bag fitted with a 5 mm strainer, which removed larger faecal particles from the suspension. The mixture was homogenized using a Stomacher 400 Circulator at 200 rpm for 30 seconds, and this process was repeated for three aliquots from each sample. A 33.3% (w/v) Histodenz solution was prepared by dissolving 1 g of Histodenz in sterile water to a final volume of 3 mL. The homogenized suspensions were pooled, layered onto the Histodenz gradients, and centrifuged at 10,000 × g for 40 minutes at 4°C. After centrifugation, 1 mL of the supernatant above the white interface was recovered as the microbial fraction **(Figure 1)**. 80µl of 0.06g/ml lysozyme was added to the samples recovered from the density gradient and incubated at 37°C for 1h at 200rpm.

Filtration of microbial suspensions is challenging because filamentous bacteria or clumped cells adhere to membranes, and so sample overloading can lead to membrane clogging (Nnadozie et al. 2015). To mitigate these issues, samples were diluted in a large volume of 1X PBS (500 mL) to improve flow. After vacuum filtration, membranes were cut into small pieces and transferred to 2 mL Lysing Matrix E tubes for cell lysis before DNA extraction. For syringe filtration, the filter was washed twice with 50 mL of 1X PBS and then backflushed with 10 mL of 1X PBS. A 1 mL aliquot of the backflushed suspension was transferred to either Lysing Matrix D or E tubes for cell lysis followed by DNA extraction.

DNA extraction was performed using the Maxwell® RSC PureFood GMO and Authentication Kit on the Maxwell® RSC48 instrument.

**Density gradients optimised to separate faecal matrix from the stool suspension**

To overcome the challenge of separating microbial cells from the fecal matrix, the stool suspension from a healthy volunteer was subjected to Histodenz density gradient centrifugation. Three single-density Histodenz solutions (33.3%, 50%, and 80% w/v) were evaluated. The 80% gradient was too dense, resulting in the fecal matrix remaining on top of the gradient and preventing clear separation from fungal cells which is used as a proxy for all microeukaryotes here. In contrast, the 33.3% and 50% gradients effectively separated the fecal matrix from the suspension, producing a distinct interface and recoverable supernatant. The 33.3% gradient provided the most complete and visually clear separation.

The bacterial loads (i.e. 16S rRNA gene amplification in qPCR) were comparable between these gradients, with mean Ct values ranging from 13.1 to 14.3 (SD > 0.5). However, a distinct difference was observed in the fungal loads (i.e. ITS amplification): the 50% gradient showed no detectable fungal signal within 40 cycles, whereas the 33.3% gradient crossed the threshold at a mean Ct of 38.6. Details of the qPCR protocol are provided in the “DNA Quantification” section.

**Optimisation of the lysozyme treatment for bacterial depletion**

The impact of lysozyme was first tested on mock communities composed of bacterial and fungal cultures **(Suppl Table 1)**. Lysozyme (0.06g/ml) demonstrated a strong antibacterial effect, reducing bacterial DNA to 18.6 ng/µl compared with the control (35.3 ± 0.61 ng/µl), while leaving fungal DNA largely unaffected (1.09 ng/µl). Therefore, lysozyme was effective in the mock community for fungal enrichments and applied further in the stool samples.

To further assess lysozyme performance, a follow-up experiment was conducted to optimise enzyme concentration. The treatment again reduced bacterial DNA (x̄ = 17.5 ± 2.43 ng/µl) relative to the reference (x̄ = 24.7 ± 6.36 ng/µl) and did not noticeably affect fungal DNA (reference x̄ = 1.3 ± 0.71 ng/µl; Lysozyme x̄ = 1.1 ± 0.08 ng/µl). However, lysozyme concentration had no or minimal impact on the degree of enrichment.

**Suppl Table 1: Composition of the mock communities**

| Bacterial | Fungal |
| --- | --- |
| *Moellerella wisconsenis* | *Candida parapsilosis* |
| *Leminorella richardii* | *Rhodotrula dairenensis* |
| *Proteus vulgaris* | *Saccharomyces cerevisae* |
| *Achromobacter xylosoxidans* | *Candida albicans* |
| *Serratia odorifera* |  |
| *Pseudomonas aruginosa* |  |
| *Serratia plymuthica* |  |
| *Staphylococcus epidermidis* |  |
| *Staphylococcus dysgalactiae* |  |
| *Enterobacter aerogenes* |  |
| *Morganella morganii* |  |
| *Salmonella enterica* |  |

**Lysing matrix tubes and their impact on fold enrichment**

Stool samples from two volunteers were used to evaluate the effectiveness of different lysing matrices on faecal samples. Each sample was first homogenised using a stomacher fitted with a 5 mm cell strainer to remove large faecal particles and produce a uniform stool suspension. The suspension was then processed sequentially through Histodenz density separation (33.3% (w/v)) to remove the faecal matrix, Lysozyme treatment (80µl of 0.06g/ml lysozyme added to 2ml of sample and incubated at 37°C for 1h at 200rpm) for bacterial depletion, and syringe filtration to enrich fungal cells, followed by lysis using five different lysing matrices for DNA extraction. Each matrix was tested in technical triplicates, and the resulting DNA was analysed by Illumina shotgun sequencing.

**DNA quantification**

DNA quantification for all samples was performed using the Qubit 4 Fluorometer and the Qubit 1X dsDNA High Sensitivity kit. The Qubit 4 Fluorometer was calibrated with standards prior to each use. For the measurements, 1 µl of DNA extract was mixed with 199 µl of Qubit™ 1X dsDNA HS working solution in a Qubit tube. The mixture was vortexed briefly and then analyzed using the Qubit 4 Fluorometer.

We quantified microbial loads during the density gradient optimisation by performing qPCR on faecal samples processed through the density gradient. SYBR™ Select Master Mix was used for qPCR, with DNA samples diluted to a concentration of 0–5 ng/µL using nuclease-free (NF) water. The reaction mixture included 10 µM primers for 16S or ITS (10 pmol/µL) (please see the primer sequences in **Suppl Table 2**), nuclease free water, and SYBR™ Select Master Mix, prepared according to the proportions detailed in **Suppl Table 3**.

**Suppl Table 2:** Sequences of primers used in qPCR reactions

| 16S | 1048F | 5'- GTGSTGCAYGGYYGTCGTCA -3' |
| --- | --- | --- |
|  | 1194R | 5'- ACGTCRTCCMCNCCTTCCTC -3' |
| ITS | ITS1F | 5’- CTTGGTCATTTAGAGGAAGTAA -3’ |
|  | ITS2 | 5’- GCTGCGTTCTTCATCGATGC- 3’ |

**Suppl Table 3:** Master mix amounts for both 16S and ITS qPCR amplification

| Component | Amount per sample (µl) **16S** | Amount per sample (µl) **ITS** |
| --- | --- | --- |
| SYBR^TM^ Select Master Mix | 3 | 3 |
| Nuclease free water | 1.6 | 1.2 |
| Primer **F** | 0.2 | 0.15 |
| Primer **R** | 0.2 | 0.15 |

In each qPCR run, nuclease free water was used as a negative control. After adding the reagents and samples, the reactions were carried out on a ViiA7 IFR E228 qPCR machine under standard settings for comparative Ct analysis. The thermal cycling program consisted of the following stages: an initial hold stage at 95°C for 3 minutes; a PCR stage of 40 cycles at 95°C for 15 seconds and 60°C for 1 minute; and a final melt curve stage, starting with 95°C for 15 seconds, followed by 60°C for 1 minute, and concluding with 95°C for 15 seconds.

All measurements were performed in triplicate, and the standard deviation of the Ct values was calculated. Measurements with a standard deviation greater than 0.5 were excluded from further analysis. For triplicates with a standard deviation of 0.5 or lower, the average Ct value was calculated and used to calculate fold enrichments.

**Preprocessing of the sequencing data and fold-enrichment calculations:**

DNA from each sample was pooled at equimolar concentrations and sequenced on an Illumina NextSeq500 using a Mid Output Flowcell (300 cycles) with a 1% PhiX spike-in. Metagenomic analysis was conducted with the in-house MATAFILER pipeline (Frioux et al. 2023). Following the protocol (Bahram et al. 2021), reads mapping to rRNA genes were identified by alignment to sequences in the SILVA 138 SSU database, and taxonomy was assigned using the LotuS2 LCA algorithm (Özkurt et al. 2022). Unprocessed samples, extracted directly without enrichment, served as references for fold-enrichment (FE) calculations, based on mean read counts across technical replicates in the genus level compositional data:

$$FE= \frac{(\sum SSU reads) - SSU Eukaryote reads}{(Ref \sum SSU reads) - Ref SSU Eukaryote reads}$$

Microeukaryotic taxa that were either unassigned at the phylum level or assigned to Arthropoda, Vertebrata, Phragmoplastophyta, Cryptomonadales, Porifera, or Schizoplasmodiida were identified and labelled as “non-target” eukaryotes. These taxa, together with taxa with a total abundance of less than 10 across all samples, were excluded from the fold-enrichment analysis to prevent inflation of fungal enrichment estimates due to false-positive taxonomic assignments to microeukaryotes.

**Suppl Table 4:** Number of reference SSU and eukaryotic SSU reads for each sample

|  |  | **Sample A** | **Sample B** | **Sample C** | **Sample D** | **Sample E** | **Sample F** |
| --- | --- | --- | --- | --- | --- | --- | --- |
| Pre-enrichment | SUM | 18097- 92780 | 36180- 49302 | 5-115 | 44371- 52578 | 26150- 29515 | 51433- 65842 |
|  | EUK_ SUM | 4-28 | 212-472 | 0 | 13-24 | 0-4 | 1-11 |
| Vacuum filtration + LysE | SUM | NA | NA | NA | NA | 10346- 19346 | 39900- 60041 |
|  | EUK_ SUM | NA | NA | NA | NA | 571-792 | 273-566 |
| Syringe filtration + LysE | SUM | 668-981 | 564- 1986 | 553-116 | 215-574 | 369-864 | 818- 1860 |
|  | EUK_ SUM | 27-176 | 3-86 | 0-29 | 12-15 | 12-20 | 22-48 |
| Syringe filtration + LysD | SUM | 2334- 5381 | 5837- 16695 | 1807- 3038 | NA | 210- 3093 | NA |
|  | EUK_ SUM | 35-59 | 121-220 | 11-38 | NA | 2-140 | NA |
